# Comparative Analysis of common alignment tools for single cell RNA sequencing

**DOI:** 10.1101/2021.02.15.430948

**Authors:** Ralf Schulze Brüning, Lukas Tombor, Marcel H. Schulz, Stefanie Dimmeler, David John

## Abstract

With the rise of single cell RNA sequencing new bioinformatic tools became available to handle specific demands, such as quantifying unique molecular identifiers and correcting cell barcodes. Here, we analysed several datasets with the most common alignment tools for scRNA-seq data. We evaluated differences in the whitelisting, gene quantification, overall performance and potential variations in clustering or detection of differentially expressed genes.

We compared the tools Cell Ranger 5, STARsolo, Kallisto and Alevin on three published datasets for human and mouse, sequenced with different versions of the 10X sequencing protocol.

Striking differences have been observed in the overall runtime of the mappers. Besides that Kallisto and Alevin showed variances in the number of valid cells and detected genes per cell. Kallisto reported the highest number of cells, however, we observed an overrepresentation of cells with low gene content and unknown celtype. Conversely, Alevin rarely reported such low content cells.

Further variations were detected in the set of expressed genes. While STARsolo, Cell Ranger 5 and Alevin released similar gene sets, Kallisto detected additional genes from the Vmn and Olfr gene family, which are likely mapping artifacts. We also observed differences in the mitochondrial content of the resulting cells when comparing a prefiltered annotation set to the full annotation set that includes pseudogenes and other biotypes.

Overall, this study provides a detailed comparison of common scRNA-seq mappers and shows their specific properties on 10X Genomics data.

**Key messages:**

- Mapping and gene quantifications are the most resource and time intensive steps during the analysis of scRNA-Seq data.
- The usage of alternative alignment tools reduces the time for analysing scRNA-Seq data.
- Different mapping strategies influence key properties of scRNA-SEQ e.g. total cell counts or genes per cell
- A better understanding of advantages and disadvantages for each mapping algorithm might improve analysis results.

## Introduction

Single cell RNA sequencing (scRNA-seq) has made great strides in the transcriptomics field as it enables differential expression analysis, clustering, cell type annotation and pseudotime analysis on a single cell level [1]. Analysis of scRNA-seq data helped to reveal new insights into cellular heterogeneity, like the altered phenotypes in circulating immune cells of patients with chronic ischemic heart diseases [2] or the transcriptional diversity of aging fibroblasts [3]. However, the analysis of scRNA-seq data is resource intensive and requires deeper knowledge of specific characteristics of each analysis tool.

The most resource intensive step during single cell NGS data analysis is the alignment of reads to a reference genome and/or transcriptome. Therefore, a common question relates to the choice of the best scRNA-seq alignment tool, that can be incorporated into a fast, reliable and reproductive analysis pipeline. Here we evaluated four popular alignment tools Cell Ranger 5, STARsolo, Alevin and Kallisto. Technological properties of these mappers are summarized in Table 1. In general, the analysis pipeline for the Chromium platform from 10X Genomics consists of Cell Ranger 5 as the standard alignment tool [4], which includes STAR [5] which was designed for bulk RNA-seq data. STAR performs a classical alignment approach by utilizing a maximal mappable seed search, thereby all possible positions of the reads can be determined. In contrast, Kallisto [6] and Alevin [7] perform an alignment-free approach, so called pseudo-alignments. The idea of alignment-free RNA-Seq quantification was introduced by Patro et al. [6] and promised much faster alignments. Here, k-mers of reads and the transcriptome are compared, and thus avoiding a comparison of each base. However, it has been shown that pseudo alignment tools have limitations in the quantification of lowly expressed genes [8].

**Table 1.**
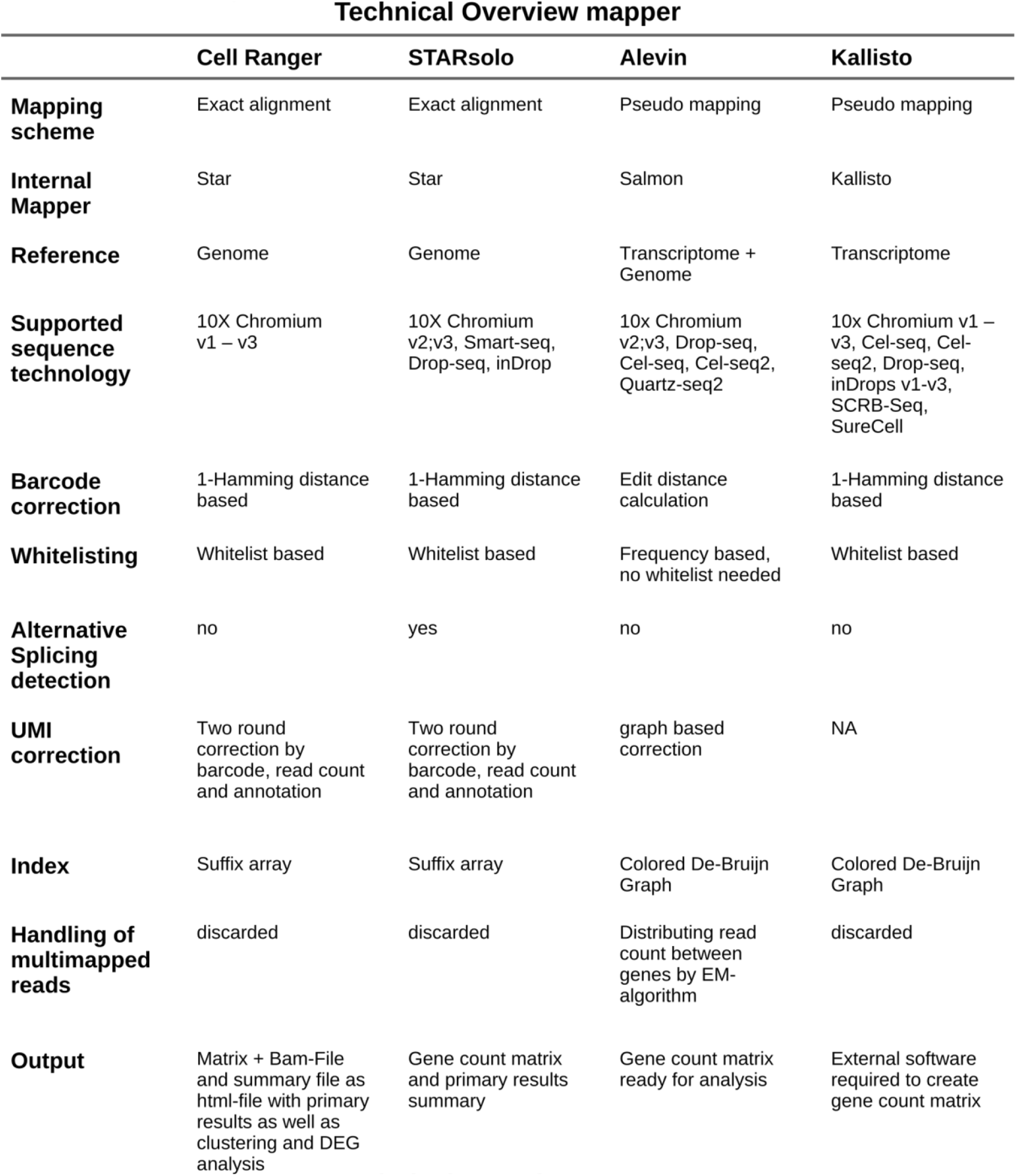
Technical overview of the most important features of each mapper.

In contrast to bulk-RNA-seq, preprocessing of scRNA-seq requires specific features. Essential features are cell calling, removing PCR duplicates and assigning reads to individual genes and cells. These features can be achieved through barcode and UMI sequences, which are sequenced along with the reads. Therefore, the correct handling of barcode and UMI sequences are crucial steps while processing scRNA-seq data. Each alignment tool applies different strategies to handle these errors. The most important step for cell calling is the correction of sequencing errors within the barcodes. Cell Ranger 5, STARsolo and Kallisto correct barcodes by comparing the sequenced barcodes to a set of all barcodes that are included in the library preparation kit, the so-called whitelist. This whitelist is provided by 10X Genomics. If no exact match of a sequenced barcode can be found in the whitelist, this barcode is replaced with the closest barcode from the whitelist, if the Hamming distance is not bigger than 1. Alevin, however, generates a putative whitelist of highly abundant barcodes that exceed a previously defined knee point. Afterwards Alevin assigns error prone barcodes to the closest barcode from the putative whitelist, while allowing an edit distance of 1.

In order to remove PCR duplicates (reads with the same mapping position, the same cell barcode) an identical unique molecular identifier (UMI) sequence is required for pooling these PCR duplicates. To correct errors in UMI sequences, Cell Ranger 5 and STARsolo group reads according to their barcode, UMI and gene annotation, while allowing 1 mismatch (MM) in the UMI sequence. As error prone UMIs are rare, they will be replaced by the higher abundant (supposedly correct) UMI. Afterwards a second round is done by grouping the barcode, corrected UMI and gene annotation. When groups differ only by their gene annotation, the group with the highest read count is kept for UMI counting. The other groups are discarded, as these reads origin from the same RNA construct but were mapped to different genes [4]. Alevin builds a UMI graph and tries to find a minimal set of transcripts for UMI deduplication [9]. In this process, similar UMIs are corrected. Kallisto applies a naive collapsing method which removes reads that originate from different molecules but contain the same UMI [10].

The third important preprocessing step of scRNA-seq data is the assignment of reads to individual genes and cells. Here the alignment tools have striking differences handling these multi mapped reads. In STARsolo, Cell Ranger 5 and Kallisto multi mapped reads are discarded when no unique mapping position can be found within the genome/transcriptome. Whereas Alevin equally divides the counts of a multi mapped read to all potential mapping positions.

Apart from the choice of the mapper, other decisions can influence the mapping results. One aspect is the choice of an appropriate annotation, which was shown to influence gene quantifications [11]. 10X Genomics recommends a filtered gene annotation that contains only a small subset that includes the biotypes protein coding, lncRNA and Immunoglobulin and T-cell receptor genes. Other biotypes e.g. pseudogenes are not included. Therefore, we were interested if a full annotation set affects the gene composition and the results of secondary analysis steps of scRNA-seq. Therefore we compared the mapping statistics of the filtered annotations to the complete (unfiltered) Ensembl annotation.

Here, we performed a benchmark of four of the most common scRNA-seq alignment tools (Cell Ranger 5, STARsolo, Alevin and Kallisto). We used different scRNA-seq data sets of mouse and human to highlight specific differences and effects on downstream analysis with a focus on clustering and cell annotation as prominent goals of droplet-based sequencing.

We are convinced that this benchmark of commonly used mappers is a valuable resource for other researchers to help them to choose the most appropriate mapper in their scRNA-seq analysis.

## Methods

### Datasets and Reference Genomes

#### 10X Drop-Seq Data

We used three publicly available data.

##### PBMC

The first data set are human Peripheral blood mononuclear cells (PBMCs) from a healthy donor provided by 10X. It was downloaded from the 10X website (https://support.10xgenomics.com/single-cell-gene-expression/datasets/3.0.2/5k_pbmc_v3). It was sequenced with the v3 chemistry of the Chromium system from 10X.

##### Cardiac

The second data set consists of 7 samples of mouse heart cells at individual timepoints (Homeostasis, 1 day, 3 days, 5 days, 7, days, 14 days, 28 days) after myocardial infarction [12]. Data was downloaded from the ArrayExpress database under the accession E-MTAB-7895. This dataset was sequenced with the v2 chemistry of the Chromium system from 10X.

##### Endothelial

The third dataset is from the mouse single cell transcriptome atlas of murine endothelial cells from 11 tissues (n=1) [13]. Data was downloaded from the ArrayExpress database under the accession E-MTAB-8077. It was sequenced with the v2 chemistry of the Chromium system from 10X. This data set could not be mapped with Cell Ranger version 4 and 5. The UMI sequence is one base shorter and the strict error handling introduced in Cell Ranger 4 could not be circumvented. Therefore, all results are based on Cell Ranger 3.

#### Gene annotation databases

Mouse and human genome and transcriptome sequences as well as gene annotations were downloaded from the Ensembl FTP server (Genome assembly GRCm38.p6 release 97 for mouse and GRCh38.p6 release 97 for human). The annotation for Cell Ranger 5 is the GENCODE version M22 for mouse and version 31 for human that match the Ensembl release 97.

In this study, we compare two annotations (filtered and unfiltered). The filtered annotation file was generated applying the *mkgtf* function for Cell Ranger v3.0.2 and *mkref* for Cell Ranger 5 according to the manual from 10X (https://support.10xgenomics.com/single-cell-gene-expression/software/release-notes/build). Therefore, the filtered annotation file contains the following features: protein coding, lncRNA and the immunoglobulin and thyroid hormone receptor genes. For the unfiltered annotation, the complete Ensembl GTF file was used without any alterations.

### Software

#### Source Code

An index of the reference genome has been built for each tool individually, using the default parameters according to the manual pages of the individual tools. The exact commands for the creation of the indices and the mapping of the data are published at https://github.com/djhn75/BenchmarkingAlignement

#### Cell filtering

Cells were filtered with the R package DropletUtils v1.6.1 [14]. All raw gene-count matrices were processed with the emptyDrops method [15]. The *emptyDrops* function applies the emptyDrops method and 50000 iterations of the Monte Carlo simulation were chosen, to avoid low resolution p-values due to a limited number of sampling rounds.

#### Downstream clustering analysis

Seurat v3.1.5 [16] was used for the downstream analysis. For all secondary analysis steps, we retained cells with a number of genes between 200 and 2500 and a mitochondrial content < 10%.

To compare the clustering we integrated the expression matrices of the samples from each mapper to remove technical noise and compare all combined samples. This was done for the Cardiac and PBMC data set. The data sets were first normalized with the *SCTransform* function. We then ranked the features with the function *SelectIntegrationFeatures* and controlled the resulting features with the function *PrepSCTIntegration*. Anchors were determined by *FindIntegrationAnchors* and afterwards used with the *IntegrateData* function. The UMAP algorithm was run on the first 15 (Cardiac) and 10 (PBMC) principal components of a PCA. To determine clusters, the *FindClusters* function was utilized with the parameter *resolution=*0.13 (Cardiac) and 0.51 (PBMC) to receive a number of clusters that is similar to the expected major cell types in the data set. The Endothelial matrices were only merged and not integrated because the resulting clustering would not yield appropriate tissue clusters due to the lack of different cell types. Yet, after merging the matrices we could obtain a similar clustering to the original study.

#### SCINA cluster comparison

To evaluate the effects of the different alignment and pseudoalignment algorithms on clustering analysis, we created an artificial “ground truth”, where we assigned each barcode to a cell type. For this task we choose SCINA v1.2 [17] as an external classification tool. The semi-supervised classification method in SCINA requires a set of known marker genes for each cell type to be classified. Marker gene sets were obtained from Skelly et. al. [18] and combined with other marker gene sets, as suggested by Tombor et.al. [19] (Suppl. Table 2). An expectation–maximization (EM) algorithm uses the marker genes to obtain a probability for each provided cell type. After the classification each cell will be assigned a cell type that shows the highest probability based on the provided marker genes. Alignments with different mappers might result in different cell classifications for each barcode. Therefore, a consensus scheme is applied to each sample to create a cell type agreement for each barcode. Consensus of a cell classification for each barcode is achieved if two or more mappers agree on a cell type.

The remaining barcodes were used as a global barcode set for SCINA. Sankey plots were generated with the R-package networkD3 (URL: https://cran.r-project.org/package=networkD3) to illustrate the representation of cell types in each Seurat cluster (Suppl. Figure 5). In addition, to convey the differences between SCINA and the seurat clusters from each mapper, a jaccard index was calculated and visualized with a heatmap (Suppl. Figure 6). UMAPs were created for each data set to illustrate the clustering between the mapper (not shown).

#### DEG analysis

For the differential gene expression (DEG) analysis each cluster from the integration in Seurat was assigned to a cell type by known marker genes for the PBMC dataset. The marker genes were obtained by the Seurat workflow for a similar 10X dataset (https://satijalab.org/seurat/v3.2/pbmc3k_tutorial.html). DEGs were then calculated by using the *FindAllMarkers* function with the Wilcoxon Rank Sum test in Seurat and all DEGs above an adjusted p-value of 0.05 were removed. Upset plots were then created with the remaining DEGs (Figure 4).

#### Additional Software

The R-package ComplexHeatmap [20] was used to create the Upset-plots (Figures 2, 4; Suppl. Figure 2).

**Figure 1.**
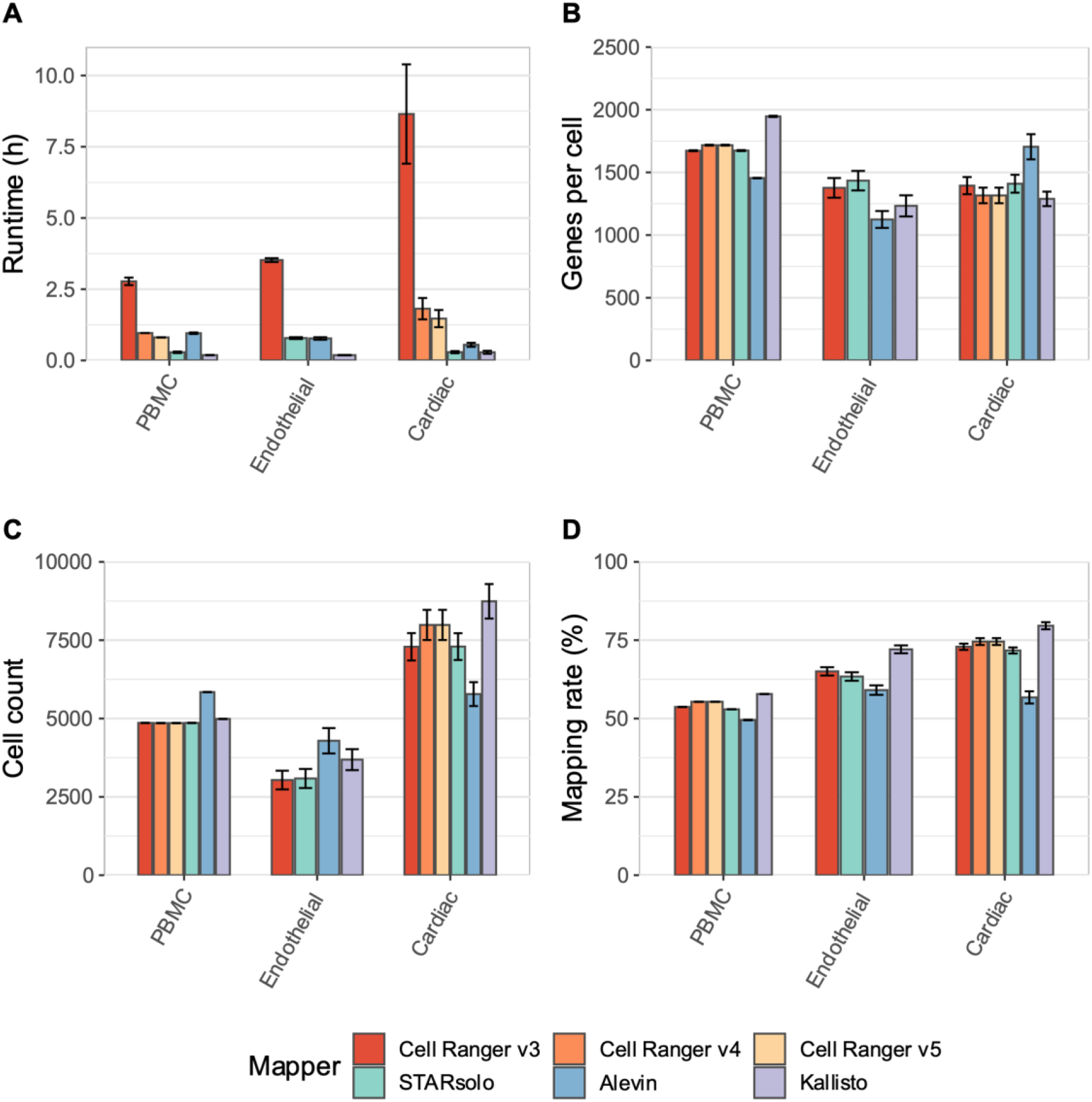
Summary of major measurements including runtime in hours (A), Genes per cell (B), cell count (C) and the mapping rate in percent (D). All bar plots show the mean of all samples with the standard error.

**Figure 2.**
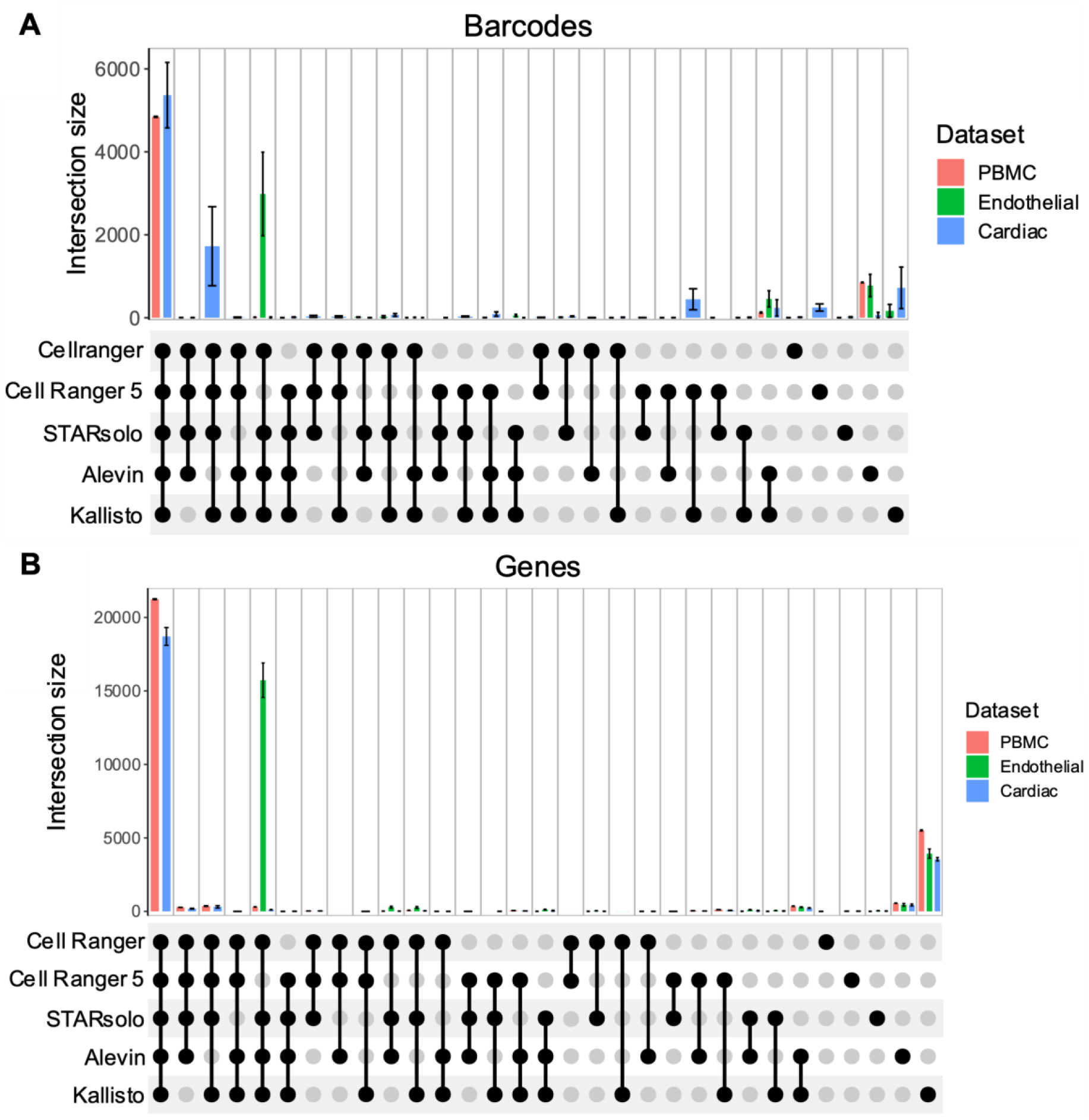
Intersection size of detected cells (A) and the detected genes (B) for each mapper. The intersection for the cells was determined by the cell-barcode. Black dots indicate the cells/barcodes that are shared among the mappers.

**Figure 3.**
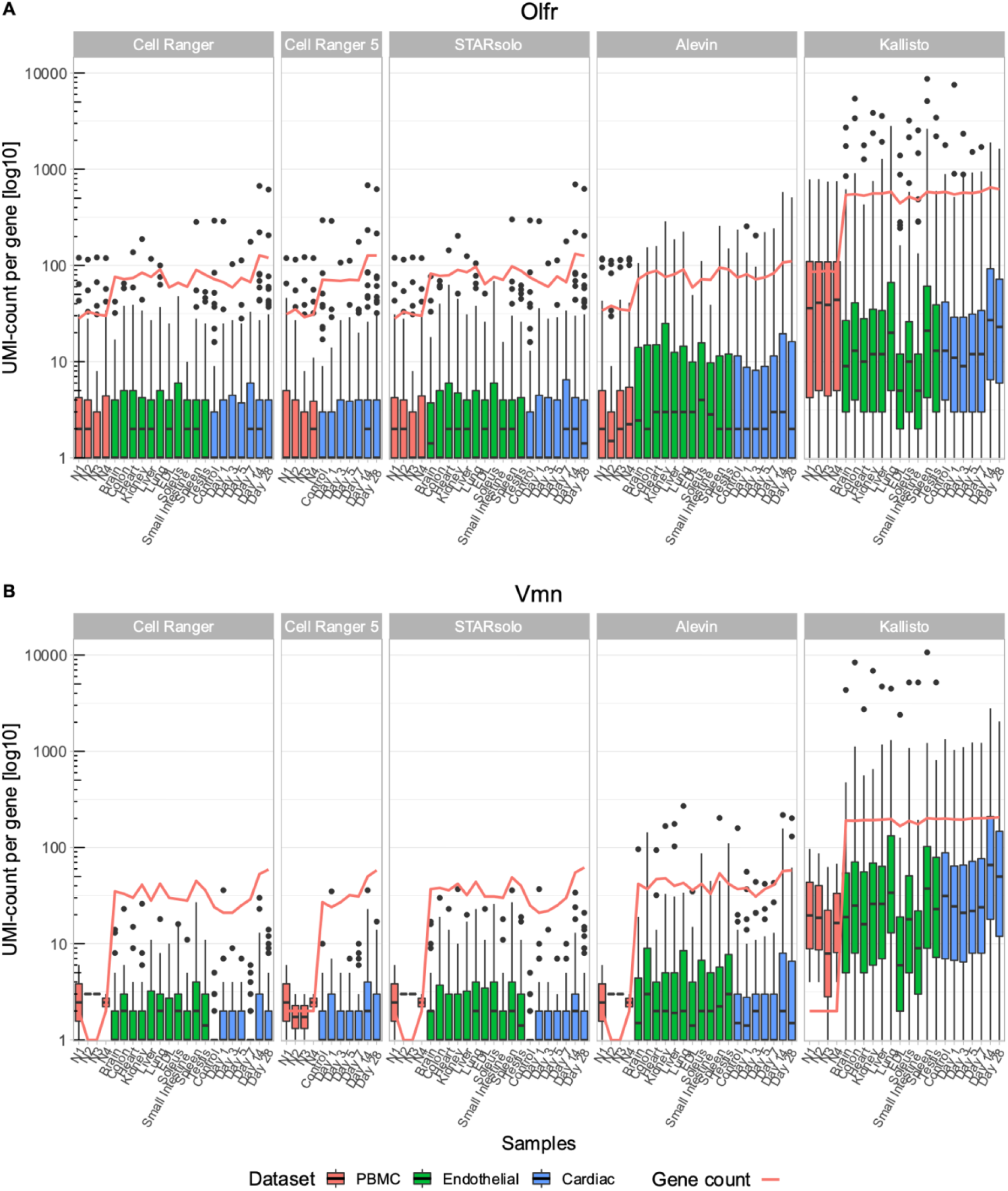
UMI counts of all detected A) Vmn (Vomeronasal receptor genes) and B) Olfr (Olfactory receptor genes) genes per mapper. The red line indicates the total number of expressed genes.

**Figure 4.**
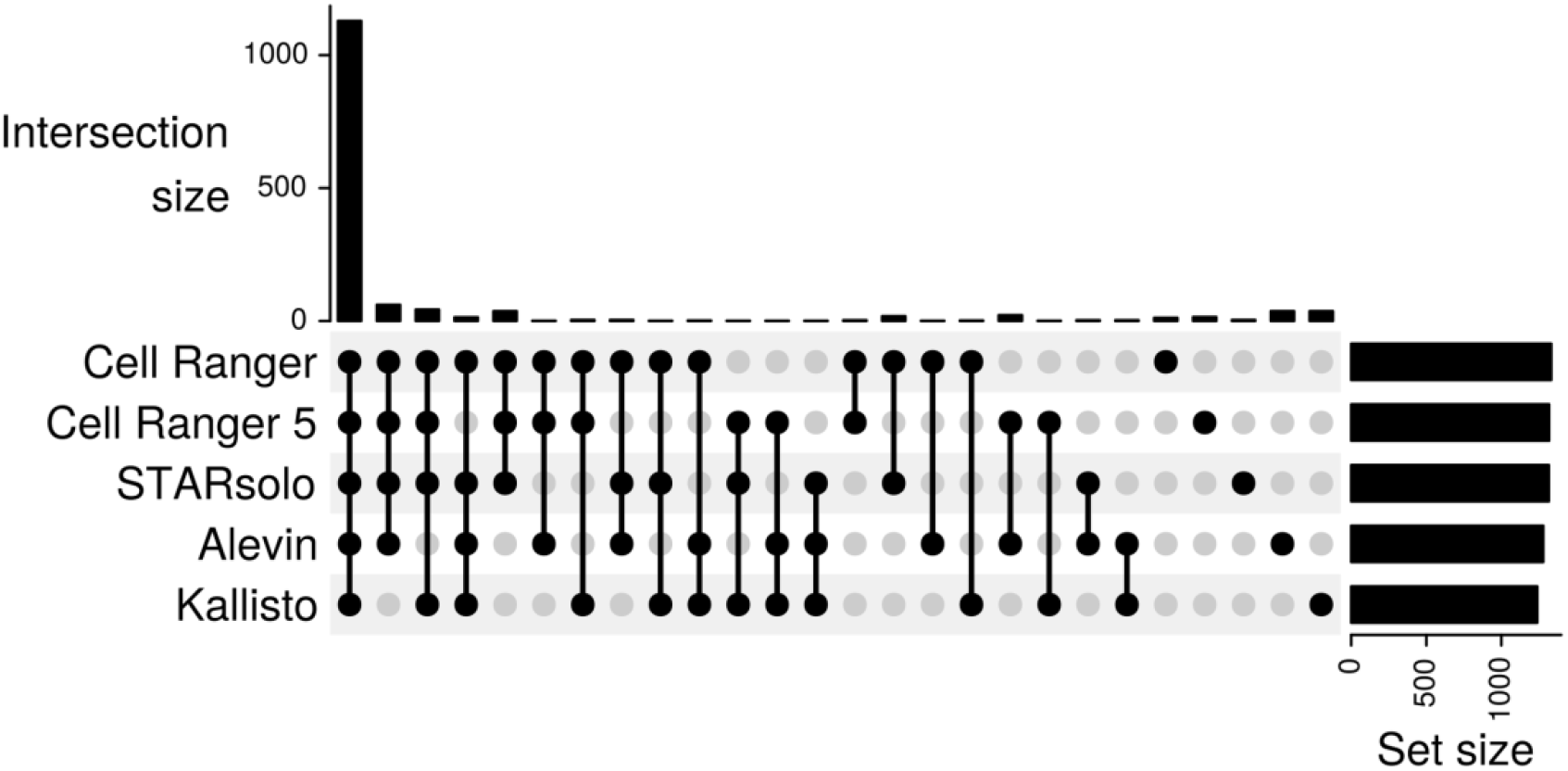
Intersection of differentially expressed genes due to the usage of different mappers in the PBMC data set.

### Hardware

All computations were executed on a workstation with Intel Xeon E5-2667 CPU and 128 GB RAM. The OS was Ubuntu 18.04 LTS.

## Results

For the comparison of the four different alignment tools Cell Ranger 5, STARsolo, Alevin and Kallisto, we analysed three representative datasets which are denoted as *PBMCs, Endothelial* and *Cardiac* (see method section for a detailed description of the datasets) in the following.

### General statistics

The overall performance and basic parameters like runtime, genes per cell, cell number and mapping rate are summarized in Figure 1. In terms of runtime STARsolo, Alevin and Kallisto clearly outperformed each version of Cell Ranger and were at least three times faster. Kallisto showed the shortest runtimes and was on average 21 times faster than Cell Ranger in version 3. With version 4 and 5 of Cell Ranger, Kallisto was 4 to 6 times faster. Additionally, Kallisto showed the highest transcriptome mapping rate whereas Alevin showed a slightly decreased mapping rate across all datasets. The cell count and the average genes per cell were similar for Cell Ranger 5 and STARsolo across all datasets. Overall Cell Ranger and STARsolo had almost identical results regarding the cell count and the genes per cell especially in Cell Ranger version 3, which is expected from the similarity of both tools. However, a slight increase of cells and a decrease of genes per cell could only be detected in version 4 and 5 in the Cardiac datasets. In contrast, Alevin and Kallisto showed different behavior for the genes per cell across the datasets. Compared to the other tools, Alevin detected more cells with less genes per cell in the PBMC and Endothelial dataset. However, it detected less cells with more genes per cell in the Cardiac dataset. More details with respect to these differences can be found in Suppl. Figure 1. In the PBMC and the Endothelial datasets, Alevin shows small peaks in the lower left corner of the density plots for UMI counts and genes per cell. These peaks represent cells which have low UMI counts. For the Cardiac dataset Alevin did not detect these cells with low UMI content, which might explain the lower cell count for this dataset. However, in the Cardiac dataset, we observed more low content cells for Kallisto. This is consistent with the finding that Kallisto detects most cells in the Cardiac dataset.

### Cell and gene identification

In 10X droplet based single cell sequencing, the individual cells are usually identified via the randomized cell barcodes, which are predefined by the whitelist. In order to determine if the different mapping tools detected identical cells, we merged the resulting cells based on their barcodes (Figure 2A). The majority of barcodes were identified by all alignment tools. However, Cell Ranger 5, STARsolo and Kallisto detected more barcodes as compared to Alevin in the Cardiac dataset. These cells had far less reads per cell compared to the cells that were detected in all mappers, as shown in the section 1 of Suppl. Figure 2 A&B. Similarly, Alevin detected unique barcodes for the PBMC and Endothelial datasets, which also had less gene content compared to the other cells detected by Alevin (panel 3 of Suppl. Figure 2 A&B). Additionally, we recognised that the majority of these barcodes are not included in the whitelist from 10X (Suppl. Table 1). Panel 4 of Suppl. Figure 2 B shows the unique barcodes for Kallisto in the Cardiac dataset, which also have less gene content than the other cells. Overall, we saw a reduced number of genes per cell for the barcodes that were only detected by one or two of the four alignment tools. By comparing the expressed genes, we could show that all alignment tools detect a similar set of genes (Figure 2B). Only Kallisto detected additional genes leading to a higher number of protein coding and lncRNA genes compared to the other tools (Suppl. Figure 3).

One gene family that occurred more frequently in Kallisto is the Vmn (Vomeronasal receptors) gene family, that is represented with higher UMI counts in the analysis performed with Kallisto (Figure 3A). Another Kallisto-enriched gene family is the Olfr (Olfactory receptor) family, which is detected with lower UMI counts compared to the Vmn family, but is still elevated compared to the other tools (Figure 3B). This leads to an increase in total gene counts for Kallisto (red line in Figure 3) and an increase of the respective biotypes (Suppl. Figure 3). The increased expression of genes from the Olfr gene family is exemplified in Suppl. Figure 4.

### Effects on downstream analysis

In order to evaluate downstream effects of the different alignment tools, we performed a semi-supervised cell type assignment with SCINA. Therefore, we used all cells that were found by more than two mappers and assigned them to a corresponding cell type based on the marker genes documented in Suppl. Table 2. Thereby, the majority of barcodes could be assigned to a specific cell type. Then we compared the clusters from each alignment tool to the assigned cell types from SCINA. Using the barcodes to identify each cell, we traced the cells from their respective clusters to the assigned cell type.

The fate from the predicted cell types to the clusters for each mapper can be observed in the sankey plots in Suppl. Figure 5. Most SCINA cell types are split into several clusters as shown in Suppl. Figure 6. In general, the clustering was similar when comparing the alignment tools. Minor differences were observed for Kallisto and Alevin. In the PBMC dataset, Kallisto showed a higher number of missing barcodes (M.b.), predominantly from monocytes. Missing barcodes are barcodes that were found in at least two of the other mappers, but not in the present one. Which means that these monocytes were not present or filtered out in Kallisto. In the Cardiac data set, the lower cell count found by Alevin leads to more barcodes associated with missing barcodes demonstrating that these cells are not detected in Alevin. The majority of these missing cells were assigned as endothelial cells. Which means that in the cardiac dataset Alevin detected only around 50% of the endothelial cells that were found with the other tools. Also the number of B-cells and granulocytes were decreased due to the lower cell counts. However, the decrease in the latter cell types could not be confirmed in the PBMC dataset.

In summary, Cell Ranger 5 and STARsolo showed the highest agreement with the predicted cell types from SCINA, which is not surprising as they use the same internal algorithm. The overlaps of Alevin and Kallisto were lower due to varying cell counts. Analysis of the differential expressed genes across the cell types of the PBMC dataset did not show major differences among the alignment tools (Figure 4).

### Comparing filtered to unfiltered annotations

The default transcriptome annotation dataset, which is recommended for Cell Ranger 5 by 10X Genomics, misses some important biotypes like pseudogenes and TEC’s, sequences that indicate protein coding genes that need to be experimentally confirmed. These differences in gene model compositions can have profound effects on the read mapping and the gene quantification as reported by Zhao et al. [11]. In order to evaluate the effects of different annotation sets on 10x scRNA-seq data, we compared the mapping statistics of the filtered annotations to the complete (unfiltered) Ensembl annotation.

Besides the increase of processed pseudogenes (Suppl. Fig. 3), the usage of the unfiltered annotation led to a decrease in mitochondrial (MT) content across all alignment tools as shown in Suppl. Fig 8A. An observation of the alignments from these genes revealed that the MT-genes were not counted in the unfiltered annotation as most of them had an equally good alignment at another position in the reference genome (multimapped reads). Suppl. Fig. 8B shows the proportion of reads for each MT-gene that are discarded because of multimappping.

It turned out that some of these MT-genes have a 100% sequence similarity with other genes (Suppl. Fig. 8C). This effect is more pronounced in mice than in humans (Suppl. Fig. 8C). Hence, a potential explanation for the reduced MT-content with the unfiltered annotation is that the mapping algorithms cannot uniquely assign a read to the MT-gene, as the read can simultaneously map to the MT-gene and the secondary sequence (Suppl. Fig. 8D). Therefore, this read is discarded.

Overall the usage of an unfiltered annotation led to a decrease of MT-content per cell. As a high MT-content is a hint for damaged or broken cells, cells with an MT-content above a certain threshold are usually filtered out from the analysis. Therefore, it will lead to an increase of total cell numbers, when applying the unfiltered annotation. Besides that, we did not see major differences in the secondary processing steps when comparing the unfiltered to the filtered annotation.

## Discussion

Since handling of scRNA-seq data is a moving target, the constant revision of new tools is important to ensure reliable results. Therefore, independent benchmarking and evaluation of uncertainties of analysis tools is of central importance [21]. Specifically for scRNA-Seq tools, comprehensive benchmarking papers are sparse [22]. Until now, only a limited number of benchmarking papers for scRNA-seq mappers were published. Du et al. [23] conducted a benchmark between STAR and Kallisto on different scRNA-seq platforms and Vieth et al. performed a pipeline comparison with simulated datasets with a vast combination of tools concentrating on imputation, normalization and calculation of differential expression [24]. Additionally, Booeshaghi and Pachter [25] performed a detailed benchmarking of Alevin and Kallisto on 10X datasets. However, an in-depth comparison of the four most common alignment tools on different 10X datasets has not been performed so far.

Our study of real 10X Genomics data sets demonstrated advantages and disadvantages of four popular scRNA-seq mappers for gene quantification in single cells and adds to the growing number of benchmarks. The tools benchmarked in this study are widely used in many labs, thus, our results are relevant for many scientists working with scRNA-seq data. All mappers have been evaluated on *in vivo* datasets as these data might reveal unexpected differences or characteristics that probably could not have been found with simulated data.

The runtime is one of the most important factors when choosing a tool, but the quality of the results is of equal importance. In our detailed analysis, we show that Cell Ranger 5 could be easily replaced with STARsolo, as they show almost identical results but STARsolo is up to 5x faster in comparison with Cell Ranger 5. The low variance in the PBMC dataset for the cell counts and genes per cell for Cell Ranger 5 and STARsolo can be explained by the predefined sample size by 10X.

Du et al. 2020 [23] reported that Kallisto was even faster than STARsolo; a finding which is consistent with our results as Kallisto had overall the shortest runtime across all mappers. However, the number of cells and the genes per cell varied across datasets for Alevin and Kallisto.

Additionally, Kallisto seems to detect genes of the Vmn and Olfr family as highly expressed in several single cell data sets, although these genes are typically not expressed in these tissues. As these gene families belong to the group of sense and smell receptors, they are expected to be expressed at lower levels or be absent in PBMCs and heart tissue and likely represent artefacts. We consistently show that these genes are overrepresented in the Kallisto results (Figure 3 and Suppl. Figure 4). As Kallisto does not perform quality filtering for UMIs this might have influenced the reported number of genes per cell as is indicated by Parekh et al [26].

Another major difference of the tested mapping tools is the handling of errors in the barcodes. We could show that Alevin often detects unique barcodes, which were not identified by the other tools. These barcodes had very low UMI content and were not listed in the 10X whitelist. It can therefore be assumed that these barcodes were poorly assigned (Suppl. Figure 2, Section 3). A possible explanation might be the usage of a putative whitelist in Alevin that was calculated prior to the mapping, instead of using the one provided by 10X.

While comparing the resulting cell clusters generated by each tool, we recognised only minor differences between the tools. Especially the clusters from Cell Rranger and STARsolo were similar. However, Kallisto detected fewer monocytes in the PBMC dataset and Alevin detected fewer endothelial cells in the cardiac dataset. Overall, we saw a much higher variance in the clustering in the cardiac dataset. This could be due to the use of an older version of the library extraction protocol (10X v2), which has short barcode and UMI sequences, or a lower sequencing quality of the Cardiac dataset.

The comparison of the complete annotation from Ensembl and the filtered annotation, as suggested by 10X, revealed that multi-mapped reads play an important role in scRNA-seq analysis. In this study, we showed that using an unfiltered annotation reduces the MT-content of cells compared to the filtered annotation. Therefore, the mitochondrial content as a way to distinguish valid cells and dead or damaged cells has to be carefully conducted as it depends on the annotation. The recommended annotation from 10X, which only contains polyadenylated genes, might lead to an overestimation of mitochondrial gene expression respectively the absence of unvalidated genes. These results suggest that there is still a need to improve the handling of multi-mapped reads in scRNA-seq data. Future mapping tools might for example consider the likelihood of a gene to be expressed in a certain cell type. This might enhance the quantification of cell type-specific genes and prevent multi-mapped reads for cell types, where a certain gene is rarely expressed. Inclusion of mapping uncertainties may be another fruitful direction.

Srivastava et al. observed that there are significant differences between methods that align against the transcriptome with quasi-mapping (e.g. Alevin) and methods that do full spliced alignments against the genome (e.g. STAR) [27]. Observed discrepancies when using the filtered annotation in our experiments often result from genes that share the same sequences, and therefore, the true alignment origin cannot be determined. The reported positions of reads contained annotated transcripts e.g. from the mitochondria and a few unprocessed pseudogenes.

In conclusion, our analysis shows that Alevin, Kallisto and STARsolo are very fast and reliable alternatives to Cell Ranger 5. They also scale to large datasets. A summary of advantages and disadvantages of each individual tool is provided in Figure 5.

**Figure 5.**
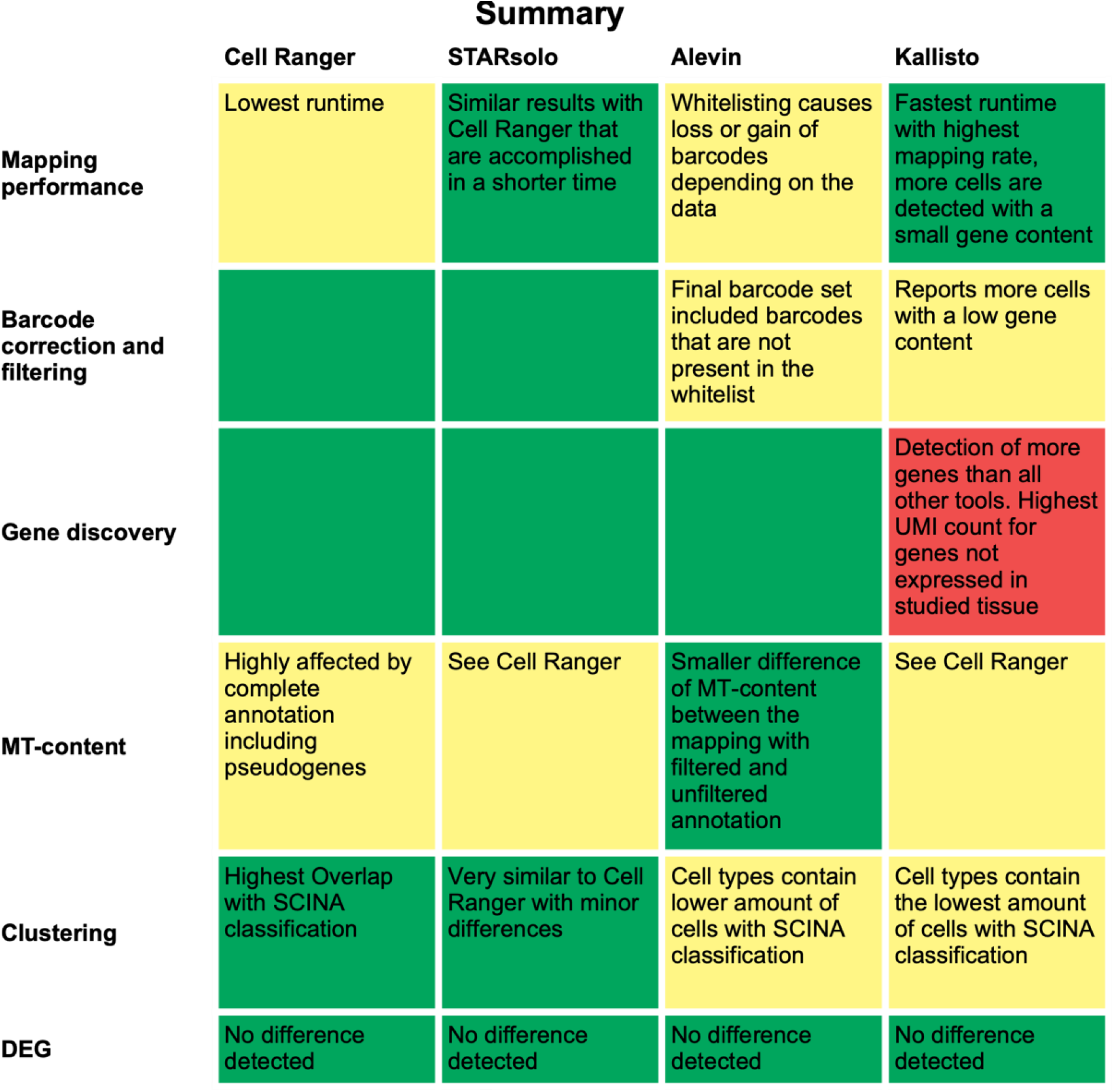
Summary of the results for each evaluated section of interest and mapper. Good results are colored in green, intermediate in yellow and poor results in red.

STARsolo should be preferred instead of Cell Ranger 5, as it is faster but otherwise performs quite comparable. If high-quality cell counts need to be obtained, Alevin appears to be the most suitable method, as average gene counts are high- and poor-quality barcodes are seldom reported. Kallisto, while reporting the highest number of barcodes, also contains many barcodes that could not be assigned to cells expected in the heart based on known marker genes.

## References

1. Wagner A, Regev A, Yosef N. Revealing the vectors of cellular identity with single-cell genomics. Nat. Biotechnol. 34:1145–1160

2. Abplanalp WT, John D, Cremer S, et al. Single-cell RNA-sequencing reveals profound changes in circulating immune cells in patients with heart failure. Cardiovasc. Res. 2021; 117:484–494

3. Vidal R, Wagner JUG, Braeuning C, et al. Transcriptional heterogeneity of fibroblasts is a hallmark of the aging heart. JCI Insight 2019; 4:

4. Zheng GXY, Terry JM, Belgrader P, et al. Massively parallel digital transcriptional profiling of single cells. Nat. Commun. 8:14049

5. Dobin A, Davis CA, Schlesinger F,et al. STAR: ultrafast universal RNA-seq aligner. Bioinformatics 2013; 29:15–21

6. Patro R, Mount SM, Kingsford C. Sailfish enables alignment-free isoform quantification from RNA-seq reads using lightweight algorithms. Nat. Biotechnol. 2014; 32:462

7. Bray NL, Pimentel H, Melsted P, et al. Near-optimal probabilistic RNA-seq quantification. Nat. Biotechnol. 34:525–527

8. Wu DC, Yao J, Ho KS, et al. Limitations of alignment-free tools in total RNA-seq quantification. BMC Genomics 19:510

9. Srivastava A, Malik L, Smith T, et al. Alevin efficiently estimates accurate gene abundances from dscRNA-seq data. Genome Biol. 2019; 20:65

10. Melsted P, Booeshaghi AS, Gao F, et al. Modular and efficient pre-processing of single-cell RNA-seq. bioRxiv 2019; 673285

11. Zhao S, Zhang B. A comprehensive evaluation of ensembl, RefSeq, and UCSC annotations in the context of RNA-seq read mapping and gene quantification. BMC Genomics 2015; 16:97

12. Forte E, Skelly DA, Chen M, et al. Dynamic Interstitial Cell Response during Myocardial Infarction Predicts Resilience to Rupture in Genetically Diverse Mice. Cell Rep. 30:3149– 3163.e6

13. Kalucka J, de Rooij LPMH, Goveia J, et al. Single-Cell Transcriptome Atlas of Murine Endothelial Cells (complete with methods). Cell 180:764–779.e20

14. Griffiths JA, Richard AC, Bach K, et al. Detection and removal of barcode swapping in single-cell RNA-seq data. Nat. Commun. 2018; 9:2667

15. Lun ATL, Riesenfeld S, Andrews T, et al. EmptyDrops: distinguishing cells from empty droplets in droplet-based single-cell RNA sequencing data. Genome Biol. 2019; 20:63

16. Stuart T, Butler A, Hoffman P, et al. Comprehensive Integration of Single-Cell Data. Cell 2019; 177:1888–1902.e21

17. Zhang Z, Luo D, Zhong X, et al. SCINA: A Semi-Supervised Subtyping Algorithm of Single Cells and Bulk Samples. Genes 10:531

18. Skelly DA, Squiers GT, McLellan MA, et al. Single-Cell Transcriptional Profiling Reveals Cellular Diversity and Intercommunication in the Mouse Heart. Cell Rep. 2018; 22:600–610

19. Tombor LS, John D, Glaser SF, et al. Single cell sequencing reveals endothelial plasticity with transient mesenchymal activation after myocardial infarction. Nat. Commun. 2021; 12:681

20. Gu Z, Eils R, Schlesner M. Complex heatmaps reveal patterns and correlations in multidimensional genomic data. Bioinformatics 2016; 32:2847–2849

21. Weber LM, Saelens W, Cannoodt R, et al. Essential guidelines for computational method benchmarking. Genome Biol. 20:125

22. Lähnemann D, Köster J, Szczurek E, et al. Eleven grand challenges in single-cell data science. Genome Biol. 2020; 21:31

23. Du Y, Huang Q, Arisdakessian C, et al. Evaluation of STAR and Kallisto on Single Cell RNA-Seq Data Alignment. G3: Genes\textbarGenomes\textbarGenetics 10:1775–1783

24. Vieth B, Parekh S, Ziegenhain C, et al. A systematic evaluation of single cell RNA-seq analysis pipelines. Nat. Commun. 2019; 10:1–11

25. Booeshaghi AS, Pachter L. Benchmarking of lightweight-mapping based single-cell RNA-seq pre-processing. bioRxiv 2021; 2021.01.25.428188

26. Parekh S, Ziegenhain C, Vieth B, et al. zUMIs - A fast and flexible pipeline to process RNA sequencing data with UMIs. Gigascience 2018; 7:

27. Srivastava A, Malik L, Sarkar H, et al. Alignment and mapping methodology influence transcript abundance estimation. Genome Biol. 2020; 21:239

