## Supplemental Figures for "Comparative Analysis of common alignment tools for single cell RNA sequencing" (from another version of this preprint: version 1, 2021-02-16)

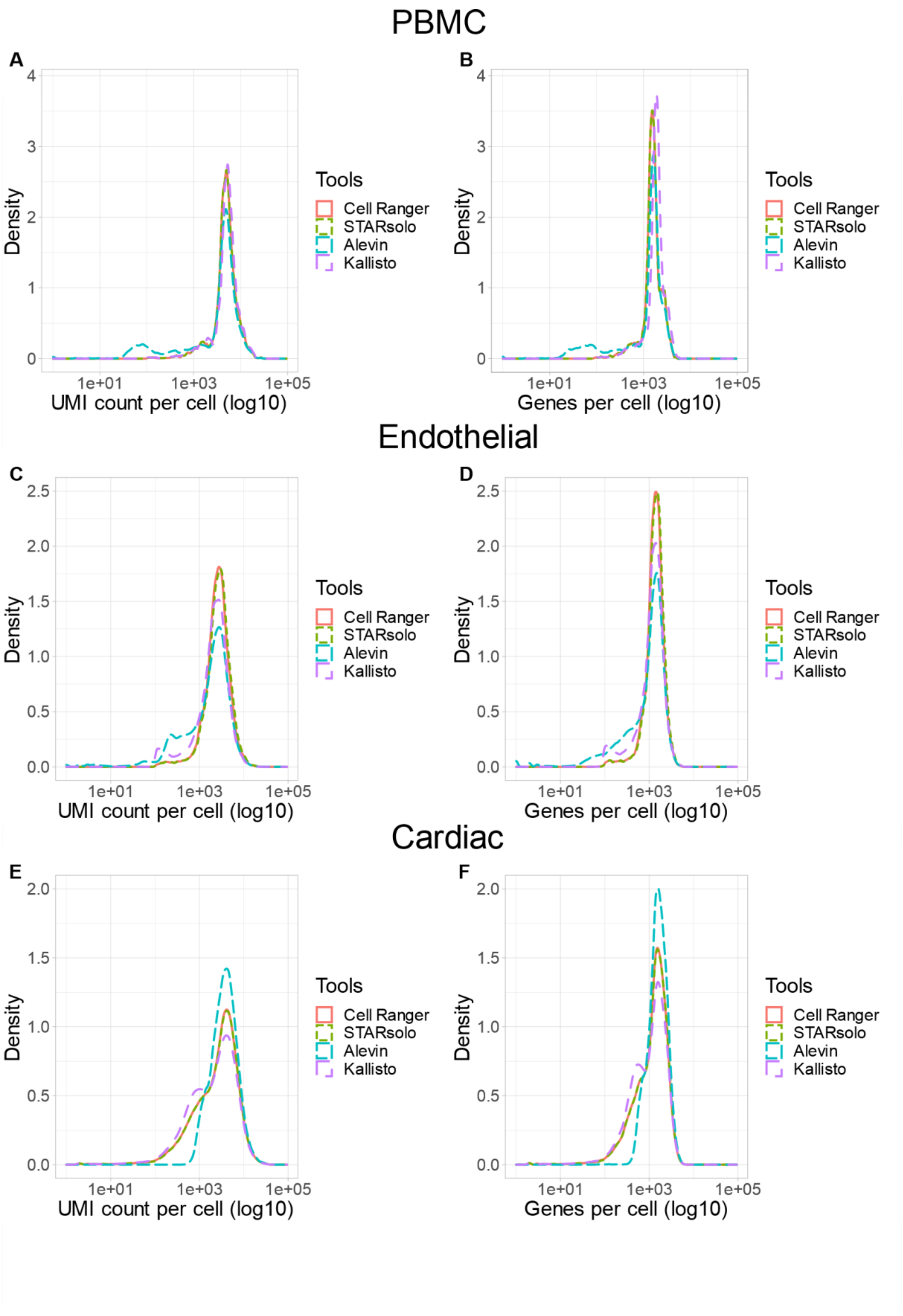


***Suppl. Figure 1*** *Distribution of UMI-counts and genes per cell for the individual data sets. Distribution is a kernel density estimate with a gaussian kernel of all samples for A, B) Rosenthal data set C, D) PBMC data set and E, F) the EC atlas data set. On the left side are the UMI counts per cell and on the right side the number of genes per cell.*

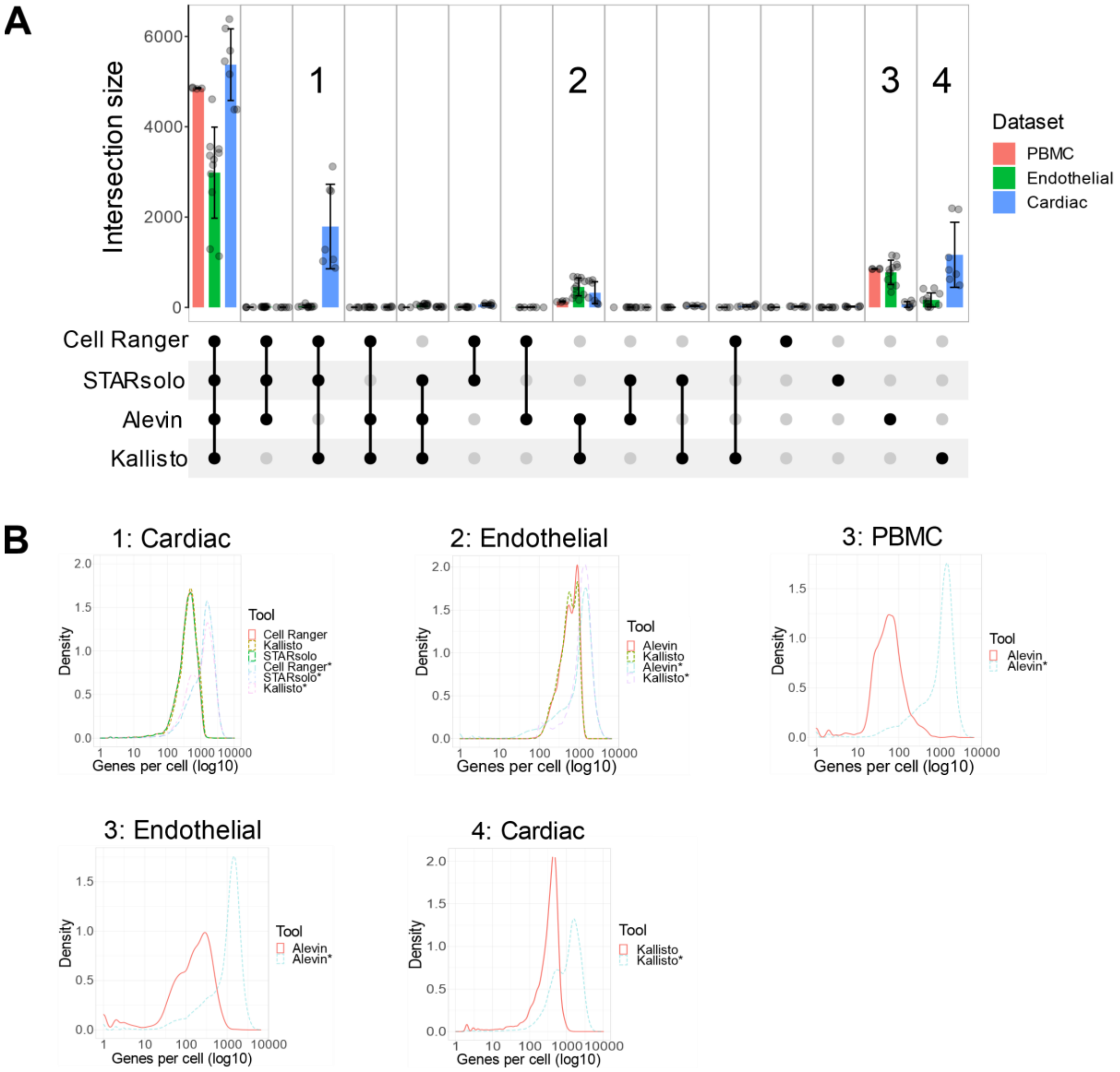


***Suppl. Figure 2*** *A.) Amount of common and unique barcodes (mean± s.e.m.) detected by the individual alignment tools. Intersections of interest are marked by numbers. B.) Gaussian distribution of genes per cells for the respective intersection and dataset from A. The distributions of the tools from the intersection (non-transparent) are compared with all detected barcodes of each tool (transparent lines (in the background); denoted with '*' in the legend)*

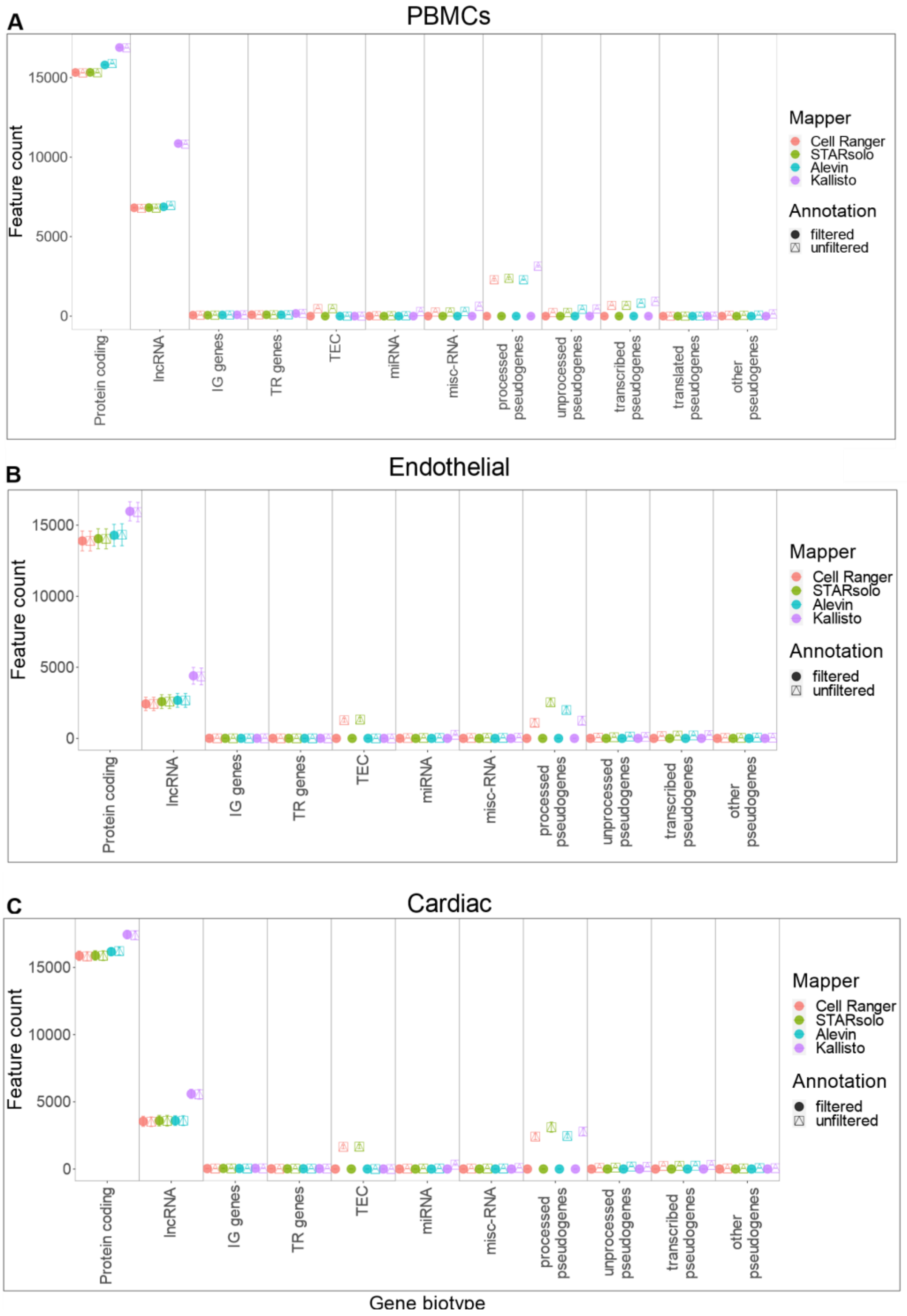


***Suppl. Figure 3*** *Number (mean+s.e.m) of biotypes per dataset with at least 1 UMI count after mapping with a filtered (solid dots) or unfiltered annotation (square-triangles). IG = Immunoglobulin genes , TR = T-cell receptor genes, TEC = Sequences that need To be Experimentally Confirmed.*

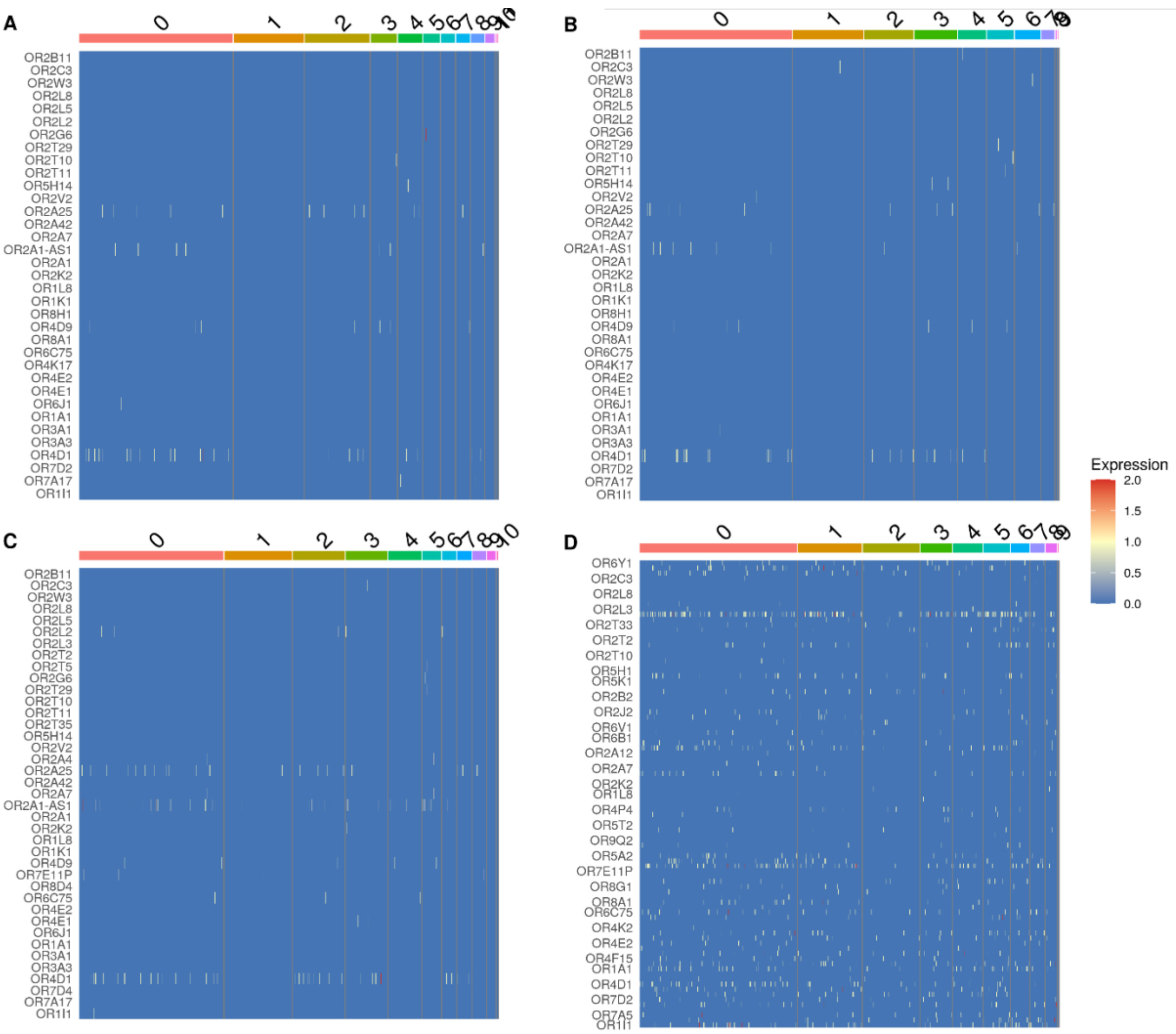


***Suppl. Figure 4*** *Expression of the OLFR gene family per cell in the PBMC data set for A) Cell Ranger, B) STARsolo, C) Alevin and D) Kallisto. Cells are sorted by clusters that are denoted by the color code above each heatmap.*

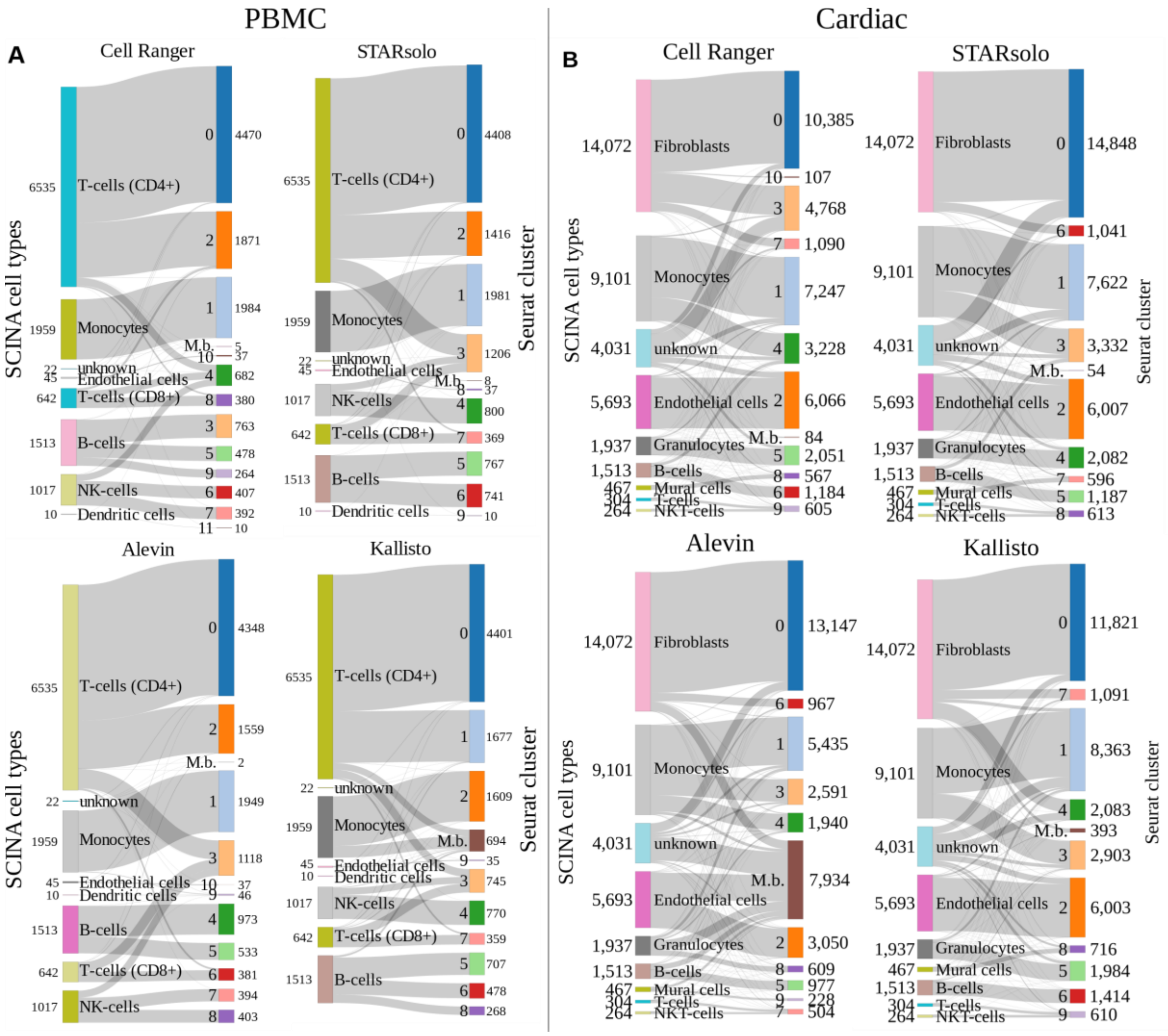


*Suppl. Figure 5 Sankey plots demonstrating the fate of each cell from SCINA cell types to the clusters obtained by Seurat. Only cells were kept if more than two mappers detected a barcode. A) represent the PBMC data set and B) the Cardiac data set. M.b. stands for missing barcodes these are barcodes that were found in at least two of the other mappers, but not in the present one.*

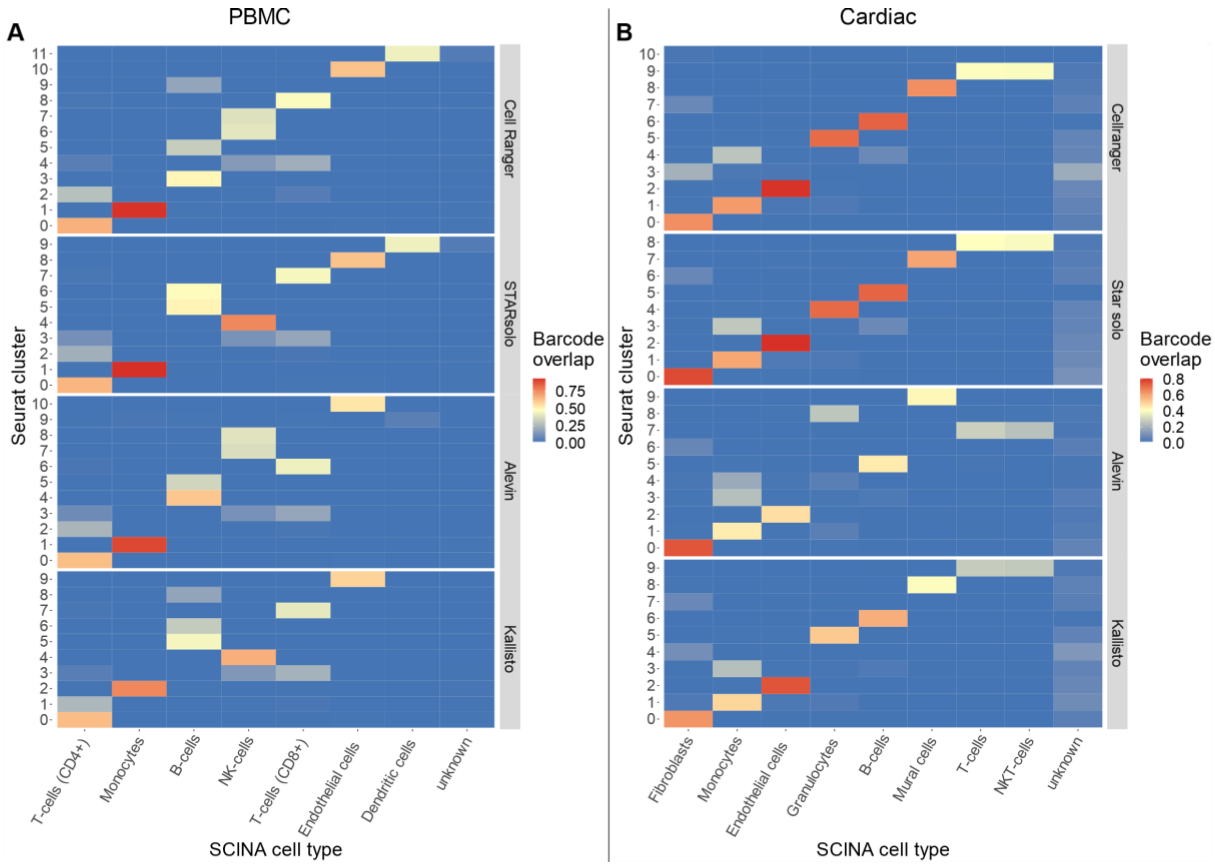


***Suppl. Figure 6*** *Heatmap showing Jaccard similarity index between Seurat cluster and SCINA cell type assignments separated by mapping tool. A) PBMC and B) Rosenthal data set. Color denotes the extent of the overlap ranging from zero (no overlap) in green, to one (full overlap) in white. Overlap is shown for each cell type.*

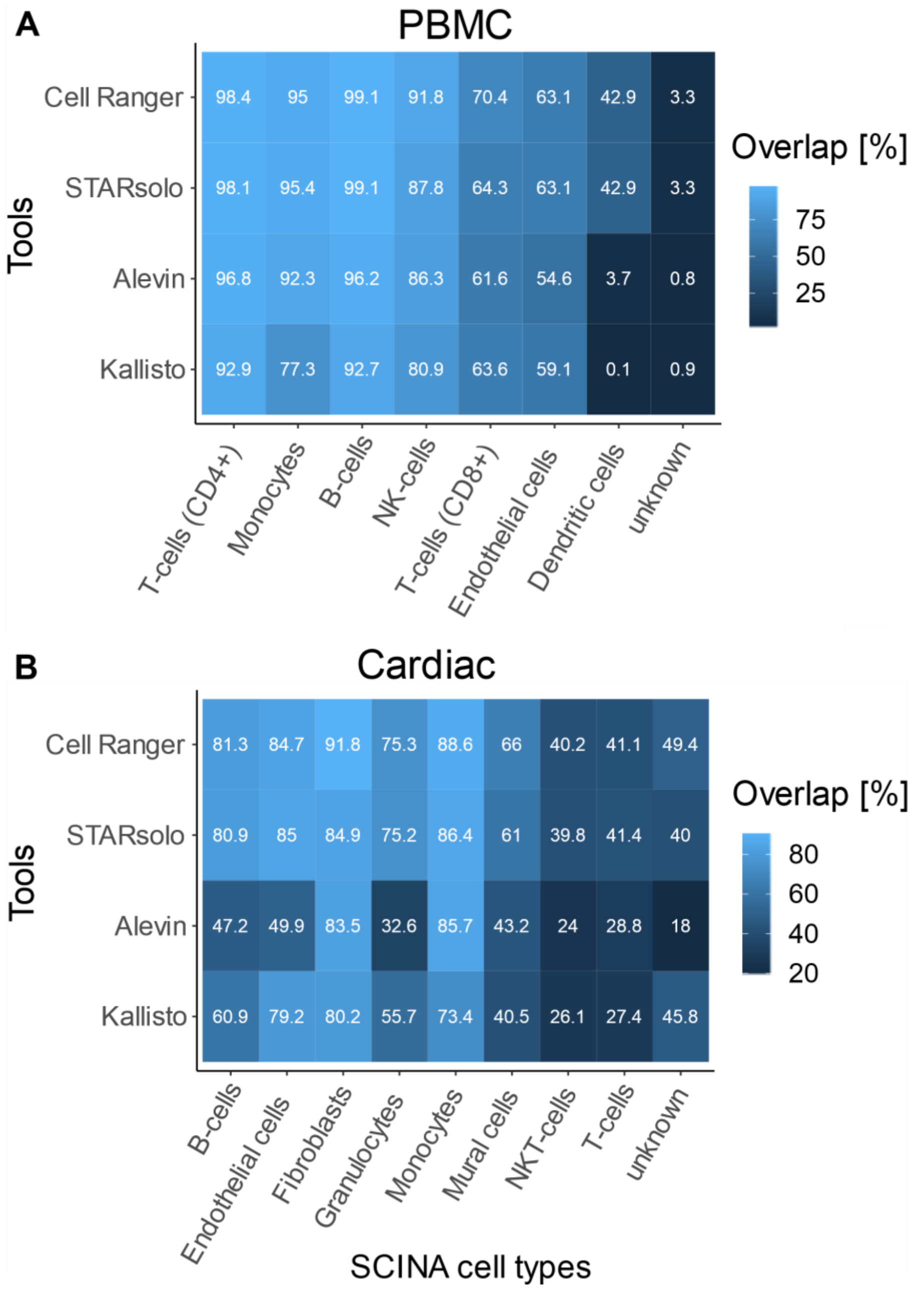


***Suppl. Figure 7*** *Average overlap of clustering from SCINA to the obtained Seurat clusters for A) the Cardiac data set and B) the PBMC data set. The overlap is calculated as the Jaccard index.*

A

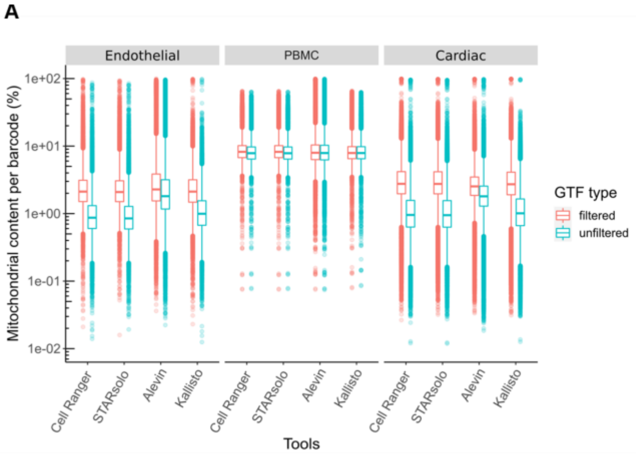


B

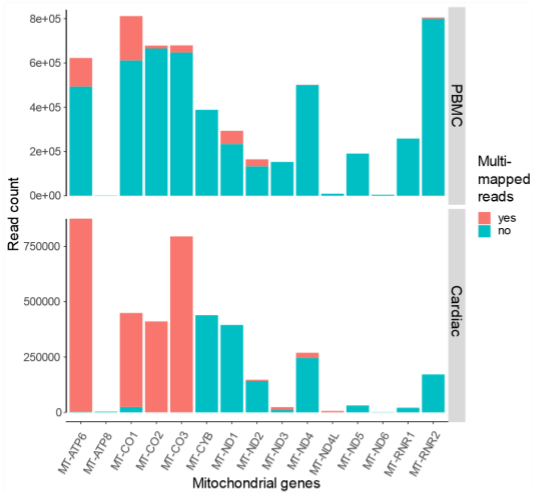


C

**Cardiac**

| Gene | Secondary mapping | Secondary mapping gene type | chr | Sequence similarity to MT gene (%) |
|---|---|---|---|---|
| Mt-Atp6 | GM10925 | unprocessed pseudogene | 1 | 100 |
| Mt-Co1 | Gm29216 | unprocessed pseudogene | 1 | 100 |
| Mt-Co2 | Gm28661 | unprocessed pseudogene | 1 | 100 |
| Mt-Co3 | Gm28437 | unprocessed pseudogene | 1 | 100 |
| Mt-Cytb | GM11410 | processed pseudogene | 4 | 92.91 |
| Mt-Nd2 | Gm13339 | unprocessed pseudogene | 2 | 96.67 |
| Mt-Nd3 | Gm28438 | unprocessed pseudogene | 1 | 99.71 |
| Mt-Nd4 | Gm28439 | unprocessed pseudogene | 1 | 100 |
| Mt-Nd4l | Gm10222 | unprocessed pseudogene | 1 | 100 |
| Mt-Nd5 | Gm11407 | processed pseudogene | 4 | 92.4 |

**PBMC**

| Gene | Secondary mapping | Secondary mapping gene type | chr | Sequence similarity to MT gene (%) |
|---|---|---|---|---|
| MT-ATP6 | MTATP6P1 | unprocessed pseudogene | 1 | 98.53 |
| MT-CO1 | MTCO1P12 | unprocessed pseudogene | 1 | 98.64 |
| MT-CO2 | MTCO2P12 | unprocessed pseudogene | 1 | 97.95 |
| MT-CO3 | MTCO3P12 | unprocessed pseudogene | 1 | 98.18 |
| MT-ND1 | MTND1P23 | unprocessed pseudogene | 1 | 99.43 |
| MT-ND2 | MTND2P28 | unprocessed pseudogene | 1 | 98.37 |
| MT-ND4 | MTND4P12 | processed pseudogene | 5 | 95.72 |

D

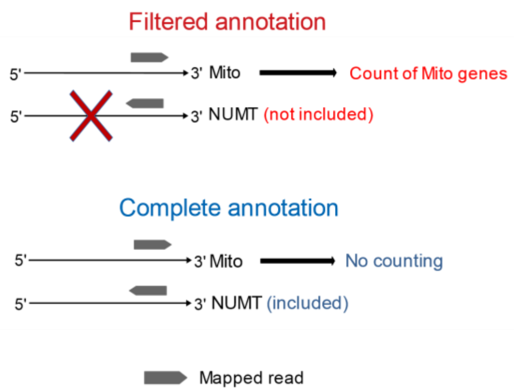


***Suppl. Figure 8*** *Difference in mitochondrial content (mt-content) of cells due to usage of a filtered and unfiltered annotation. A) MT-content of cells separated by filtered and unfiltered annotation. B) Reads mapped to the mitochondrial genes for the PBMC and Rosenthal data set with unfiltered annotation. Red indicates proportion of reads that are removed due to multimapping. C) Mitochondrial genes and its closest pseudogene when the mappers reported the secondary mapping position along with the sequence similarity to the MT gene. D) Example of the mapping process of a read from a MT gene with a filtered/unfiltered annotation. As the filtered annotation does not include potential NUMT's, the read is uniquely mapped to the MT gene. Whereas the complete set contains NUMT's and therefore the read cannot be uniquely mapped to the MT genes (multimapped) and therefore is discarded from counting.*
